# The pseudogene SURFIN 4.1 is vital for merozoite formation in blood stage *P. falciparum*

**DOI:** 10.1101/562124

**Authors:** Tatiane Macedo-Silva, Rosana Beatriz Duque Araujo, Gerhard Wunderlich

**Author notes:** correspondence to: Gerhard Wunderlich.

## Abstract

The *surf* gene family of the human malaria parasite *Plasmodium falciparum* encodes for antigens with largely unknown functions. Three of the ten *surf* genes found in the *P. falciparum* 3D7 genome are annotated as pseudogenes, and one of these – *surf*4.1 (PF3D7_0402200) - was continuously transcribed in *P. falciparum* 3D7 blood stage forms. GFP-tagging revealed that despite several stop codons a full-length protein was expressed, which localized to developing merozoites. Analysis of cDNAs showed that no specific editing occurred pointing to readthrough of stop codons during translation. Intriguingly, attempts to generate parasite lines containing an additional artificial stop codon failed. Transcript knockdown revealed that *surf*4.1 is essential for merozoite formation in late trophozoite/schizont stages while DNA replication seemed not to be influenced. SURFIN4.1 is the first example of a plasmodial multigene family member of which a knockout is deleterious and may pose as a novel target for anti-malarial therapy.

## Introduction

Malaria is still one of major infectious diseases in the world and despite huge advances in the control of the disease, a recent stalling in the decrease of cases is alarming (World Health Organization, 2018). Malaria is caused by apicomplexan protozoans of the genus *Plasmodium* and the most lethal species for humans is *Plasmodium falciparum*. It is estimated that only in 2017 435,000 people died of malaria and 219 million new cases were reported. Most of the victims occur in sub-Saharan Africa and are children younger than 5 years (World Health Organization, 2018).

In the vertebrate host, parasites invade red blood cells and go through intracellular multiplication by a process termed schizogony which culminates in the lysis of the infected red blood cell (IRBC) and the liberation of up to 32 merozoites which instantly reinvade erythrocytes. The release of toxic byproducts and cytoadherence of infected red blood cell to other uninfected red blood cells or receptors in deep venules are the main triggers for the severe outcomes of *P. falciparum* malaria in individuals without adequate immunity against infection (revised in (van der Heyde et al., 2006; Milner, 2018)). In the parasite genome, a number of multigene families are present, and in many cases, these encode variant proteins which are involved in host immune response evasion, and also in pathogenic processes, such as PfEMP1 (Baruch et al., 1995; Smith et al., 1995; Su et al., 1995), RIFINs (Fernandez et al., 1999), STEVOR (Cheng et al., 1998) and others (revised in (Wahlgren et al., 2017)).

Another, smaller gene family comprises *surf* genes (surface-associated interspersed family) (Winter et al., 2005), which appear in different numbers in the *P. falciparum* strains sequenced so far. In contrast to *rif* and *var* genes which encode RIFINs and PfEMP1 antigens, respectively, *surf* genes are apparently more conserved between different strains and probably do not undergo frequent ectopic recombination as shown for *var* genes (Freitas-Junior et al., 2000). Additionally, in *Plasmodium vivax*, the PvSTP gene family (del Portillo et al., 2001) shows significant similarities to *surf* genes, considering the domain structure of a small variant ectodomain-encoding part and a larger, tryptophan-rich regions-encoding domain (Winter et al., 2005). SURFINs are large antigens with more than 200 kDa. In *P. falciparum* strain 3D7, ten genes encode *surf* genes and three of these are annotated as pseudogenes. The presence of a modified export motif (PEXEL/VTS (Hiller et al., 2004; Marti et al., 2004)) in the N-terminal region of SURFINs in some of the alleles suggested that SURFINs are exported to the IRBC surface and first results analyzing the allele SURFIN 4.2 (Winter et al., 2005) revealed that this protein was present on merozoites and partially on IRBCs, colocalizing with PfEMP1. In a recent study, it was observed that SURFIN 4.2 from the FCR3 line forms complexes with RON4 (rhoptry neck protein 4) and GLURP (Glutamate-rich protein). Also, antibody-mediated inhibition of RBC invasion was documented, suggesting a role in parasitophorous vacuole formation. Intriguingly, *surf*4.2 from the CS2 line can be knocked out without any growth defect (Maier et al., 2008) and recent genome-wide mutational analysis indicated that *surf*4.2 may be dispensable with a small fitness cost upon insertion-mutation (Zhang et al., 2018). Few data are available regarding other *surf* alleles. Mphande and colleagues confirmed the localization of SURFIN 4.1 on the merozoite surface but not on the surface of IRBC (Mphande et al., 2008). The same authors also detected that *surf*4.1 occurred in six copies in the FCR3 strain, while only one copy was present in the strain 3D7 and the Brazilian isolate 7G8. Later, Ochola and colleagues detected in a genome-wide approach that *surf*4.1 is under selective pressure in circulating field isolates meaning that the protein is a target of the naturally acquired immune response (Ochola et al., 2010). A similar analysis examining *surf*4.1 ectodomain-encoding sequences from Thai isolates showed diversifying selection, although the frequency of allele distribution was stable over years in the analyzed isolates (Xangsayarath et al., 2012). More recently, Gitaka and authors analyzed the surprisingly high occurrence of frameshift mutations in the *surf*4.1 open reading frame (ORF) in field samples and hypothesized that this may lead to truncated versions of the protein (Gitaka et al., 2017). Similar to *surf*4.2, *surf*4.1 was deemed dispensable in genome-wide insertion-mutation assays (Zhang et al., 2018), although insertion was only observed near the 3’ end of the *surf*4.1 open reading frame. During an effort to analyze the mode of transcription of the *surf* gene family and eventually observe transcriptional switching, we generated NF54-based parasite lines (isogenic with strain 3D7) with either a green fluorescent protein-hemagglutinin tag or a degrading domain, variant DD24 (de Azevedo et al., 2012), fused to tagged to the SURFIN4.1 polypeptide. Although there are a number of stop codons in the NF54/3D7 *surf*4.1-ORF, green fluorescence was readily observed and in western blots, a full-length protein was detected (Macedo-Silva et al., 2017). Also, all parasites were positive for GFP presence, ruling out parasites with only truncated versions of the protein. In the same study, significant transcriptional activity was found from the *surf*4.1 locus, and other loci were only sporadically activated during multiple reinvasions. Further, although an 80% protein knockdown of SURFIN4.1 led to no discernible growth defect, a significant increase in the steady-state levels of the *surf*4.1 transcript was observed during knockdown which returned to normal values when the protein was reestablished (Macedo-Silva et al., 2017). This phenomenon may be interpreted that the parasite counterbalances the lack of viable protein by increasing transcription. If so, a further decrease of SURFIN 4.1 may be deleterious for parasite survival. Coincident with this, a successful knockout of *surf*4.1, if attempted, was never reported. Together with all previous observations, these results point to a specific role of *surf*4.1 at least in the NF54 or 3D7 line. Here, we addressed the biological function of *surf*4.1 by introducing an efficient knockdown at the transcriptional level and provide evidence that *surf*4.1 is indeed an essential protein, probably involved in merozoite formation during red blood cell schizogony.

## Material and Methods

### Plasmid constructs

DNA fragments encoding parts of the *surf*4.1 gene (PlasmoDB ID PF3D7_0402200) were amplified by PCR (See Supplementary Table 1 for sequences) using Elongase proofreading enzyme (Invitrogen). The amplicons were cloned in pGEM T-easy vectors (Promega) and sequenced. The SURFIN4.1-3′ ORF encoding fragment was excised using Bgl2 and Pst1 and transferred to a modified pRESA-GFP-HA vector (de Azevedo et al., 2012) digested with the same enzymes. This vector had the glmS element inserted downstream of the stop codon, resulting in pS4GFPHAglmS. The plasmid pS4GFPHAglmsTK was cloned using two homology regions for double crossover recombination. For this, the *surf*4.1 3’ ORF fragment was excised together with the GFP-HAglmS encoding region plus its terminator via BglII and EcoRV and inserted in the same site in pHHTK (Duraisingh et al., 2002). Afterwards, the 3’UTR was amplified, cloned in pGEM T easy, sequenced and transferred to the pHHTK vector using NcoI and ClaI. The plasmid pS4(TAA)GFPHAglmsTK was constructed similarly with the exception that a fusion-PCR was performed to introduce a single nucleotide exchange (T→A) at position 5663. The fused, mutated 3’-ORF fragment was transferred into pS4GFPHAglmS (via BglII and NheI), resulting in pS4(TAA)GFPHAglmS, from which it was transferred via BglII/EcoRV to pHHTK already containing the 3’UTR homology region to create pS4(TAA)GFPHAglmsTK. The construction of pS4KO for 5’ and 3’ ORF was done using the pHHTK backbone inserting the respective fragments into the BglII/EcoRI and Spe1/NcoI restriction sites. Recombinant plasmids were grown to high quantities using the Maxiprep protocol (Sambrook, 1991) and used for transfections. All oligonucleotides used in the amplification and cloning steps are informed in Supplementary Table 1.

### Parasite culture and transfection

*Plasmodium falciparum* lineage NF54 (Walliker et al., 1987), kindly provided by Mats Wahlgren (Karolinska Institutet, Sweden) was used throughout the experiments. Blood stage parasites were maintained in RPMI supplemented with 0,23% NaHCO_3_, 0.5% Albumax 1 (Gibco, Rockville MD) and human B+ erythrocytes in a defined gas mixture (90% N_2_, 5% O_2_ and 5% CO_2_) (Trager and Jensen, 1976). The synchronization of parasites was done by plasmagel flotation (Lelievre et al., 2005) of mature trophozoites followed by sorbitol lysis (Lambros and Vanderberg, 1979) of ring stage forms. Transfection of schizont stage parasites was done using the protocol published by Hasenkamp and colleagues (Hasenkamp et al., 2012). Transfected parasites were grown using 2.5 nM WR99210 (a gift from Jacobus Inc., USA). For the integration via single crossover recombination, transfected parasite lines were cultivated for 14-20 days without WR99210, after which the drug was added again. Normally, after three cycles locus-integrated parasite lines were obtained. These were cloned by limiting dilution. Experiments were done in biological duplicates or triplicates. For the integration via double recombination, following the outgrowth of transfected parasites, these were cultivated for 15 days without WR99210 and after re-adding the drug, the parasitemias were adjusted to 1%. Then, 2.2 µM ganciclovir was added into the culture for 6 days, followed by 4 days at ganciclovir concentrations of 4.4 µM and 4 days at 8.8 µM. Afterwards, genomic DNA of parasites was extracted to check for successful integration and elimination of episomal plasmid forms.

### Knockdown assay

To knockdown protein expression in SURFIN4.1GFPHAglmS parasites, D-glucosamine 6-phosphate (Sigma-Aldrich G5509) was added to a final concentration of 2.2 mM to highly synchronous ring stage parasites (6 h post reinvasion, hpi). Parasites were kept in culture until the schizont phase (38-46 hpi), monitored by microscopy. For protein extraction, pelleted red blood cells were treated with 0.1% saponin in PBS (supplemented with Complete Protease Cocktail Inhibitors (Roche)) to remove hemoglobin. Proteins were electrophoresed on standard discontinuous SDS-polyacrylamide gels, transferred to Hybond C membranes (Amersham) and analyzed as described below. The same saponin lysis process was done for RNA extraction applying 1 ml of Trizol reagent (Life Technologies) to pre-purified pelleted parasites and subsequent storage at −80°C until use.

### Sequencing of genomic DNA and complementary DNA

Total genomic DNA of parasites was extracted using the Wizard® Genomic DNA Purification Kit (Promega) following the manufacturer’s instructions. cDNA was synthesized from total RNA purified from Trizol-treated samples (Life Technologies) following the manufacturer’s instructions (see below). From resulting cDNA and gDNAs, fragments of *surf*4.1 spanning the sequence of interest were amplified by PCR using Elongase (Invitrogen) with primers specified in Supplementary Table 1 and cloned into vector pGEM-T Easy (Promega). A number of resulting clones was Sanger-sequenced and analyzed using the BLAST tool at PlasmoDB (www.plasmodb.org). Sequence alignments were also done using ClustalX2.1 (Larkin et al., 2007).

### Immunoprecipitation (IP) and mass spectrometry (MS)

For IP, parasite protein extracts were prepared as described above. The Pierce Co-IP kit (MACS® MicroBeads) was used following the manufacturer’s instructions. In brief, anti-GFP coupled to beads was added to parasite extracts from late schizonts and incubated for 2 h at 4°C. Afterwards, the samples were passed through a magnetic column followed by three washes and two elution steps. Immunoprecipitated fractions were visualized by SDS-PAGE and Coomassie blue staining. MS analyses were performed on eluted proteins on an LTQ-Orbitrap Velos ETD (Thermo) coupled with Easy nanoLC II (Thermo). The peptides were separated on a C18 RP column on a 115 min gradient. The instrumental conditions were checked using 100 fmol of a tryptic digest of BSA as standard. Peptides were identified using the ProteinDiscovery software tool (Thermo) against a *P. falciparum* databank from Uniprot.

### Realtime qPCR

For *surf* transcript quantification, 9 oligo pairs corresponding to the 3D7 *surf* genes available in PlasmoDB (version 8) were used (Supplementary Table 1). Notably, *surf* genes 3 and 8 are identical. Whole RNA was purified from synchronized stages (ring stage, directly after sorbitol treatment, trophozoite stage, 20 h after sorbitol treatment, and schizont stage, 30 h after sorbitol treatment) by the Trizol protocol and dissolved in pure water. RNAs were then treated with DNAse1 (Fermentas) and cDNA synthesis was done using RevertAid reverse transcriptase (Fermentas) using random oligos as published earlier (Gölnitz et al., 2008). As an endogenous control transcript, the plasmodial serine tRNA ligase transcript (PlasmoDB PF3D7_0717700), was used.

### Immunoblotting

Whole parasite protein extracts were prepared from saponin-lysed IRBCs as described in Methods in Malaria Research (Ljungström et al., 2008). Proteins were loaded on standard discontinuous SDS-polyacrylamide gels and transferred to Hybond C membranes (Amersham). After blocking with 5% skimmed milk in 1xPBS/0.1% Tween20, HA-tagged proteins were recognized using a murine antiHA antibody (Sigma-Aldrich) and then an antiMouse IgG-peroxydase antibody (KPL). Blots were exhaustively washed with PBS/Tween between incubations and finally incubated with ECL substrate (GE). As a loading control, a murine anti-PTEX150 (generously provided by Dr. Mauro Azevedo) or an anti-Histone 3 antibody (Cell Signaling Inc.) was used. Chemoluminescent signals were captured in an ImageQuant (GE) apparatus and/or X-ray films and intensities were quantified using ImageJ software (NIH). The obtained values were normalized using the PTEX150 or Histone 3 signals.

### Microscopy for GFP Fluorescence and Immunofluorescence

To analyze GFP fluorescence, synchronous parasites were collected immediately before use. After incubation with DAPI (5 µg/ml) for nuclear DNA staining, 20 µl of erythrocytes were placed on a glass slide, covered with a coverslip, and analyzed on a Zeiss Observer Axio Imager M2 microscope with 1000x magnification.

For immunofluorescence, the samples were collected and prepared using the protocol published by Tonkin and colleagues (Tonkin et al., 2004).

### Southern Blot

10µg genomic DNA of lineages NF54 (control) and NF54::pS4GFPHAglms parasites, isolated using the Wizard® Genomic DNA Purification Kit (Promega), and 25 ng of plasmid DNA, were digested using BamHI (Fermentas/Thermo). The probe was amplified by PCR with digoxigenin-dUTP from the DIG High Prime DNA Labeling and Detection Starter Kit I (Roche Diagnostics), using the glms sequence as a template, amplified with oligos 5′-TAATTATAGCGCCCGAACTAAGC-3′ and 5′-AGATCATGTGATTTCTCTTTG-3′. Hybridization and visualization were done following the manufacturer’s instructions (Roche), using Hybond N membranes (Amersham/GE Healthcare) at a hybridization temperature of 45°C.

## Results

### The genomic sequence and the reverse translated message of *surf*4.1 have stop codons

In order to confirm if the predicted stop codons were indeed present in the NF54 parasite line used throughout our experiments, a number of plasmid clones containing amplicons from a *surf*4.1 region that contain the splicing site and stop codons were sequenced. The same was done for plasmid clones made from cDNA-originated amplicons. Of note, a proofreading enzyme was included in PCRs in order to avoid sequence artifacts. The used NF54 parasite line constitutively transcribes *surf*4.1 (Supplementary Figure 1).

Translating from the predicted start ATG, we observed that a first stop codon occurred at nucleotides 2488-2490. The only difference found was a six-thymidine deletion (2447-2452) inside a highly repetitive region in the intron, which may indicate that the used NF54 strain differs at this point from the deposited 3D7 strain sequence in PlasmoDB (data not shown). When analyzing the sequences from cDNA-derived amplicons, the splicing of the predicted intron at nt 2371 to nt 2464 was observed, however, another predicted non-canonical two-base-pair intron (2530-2531) was not spliced out (Supplementary Figure 2). When starting from the predicted translation start ATG, this insertion leads to a stop codon (TGA) at this point and several others from this point on. Moreover, all other possible ORFs resulted in stop codons. This means that the parasite must employ read-through of stop codons when translating the *surf*4.1 into protein.

### Tagging of *surf*4.1 with *gfp* leads to fluorescent forms of full length SURFIN4.1-GFP-HA

To examine the presence of protein SURFIN4.1-tagged GFP in blood stage parasites, we created an NF54 parasite line which had its *surf*4.1 gene genetically tagged with a GFP-HA tag (Figure 1). After drug on/off-cycling and cloning by limiting dilution, the transfectant parasite lines containing an integrated version of the plasmid pSURF4-GFP-HA were PCR tested and no amplification product was obtained with an oligo pair which detects episomal forms (Figure 1B). When blotting a whole parasite extract of the transfectant line, SURFIN4.1-GFP-HA was detected in its correct size using an antiHA antibody in Western blots (Figure 1C). The parasite line containing an integrated version of the plasmid showed green fluorescence in late schizonts in cytometry analyses (data not shown) as well as in fluorescence microscopy (Figure 1D). No increase in abnormal parasite forms or extension or decrease of the blood stage cycle duration was noted. This reinforces that a read-through of stop codons must occur during translation of the *surf*4.1-gfp-ha transcript.

**Figure 1:**
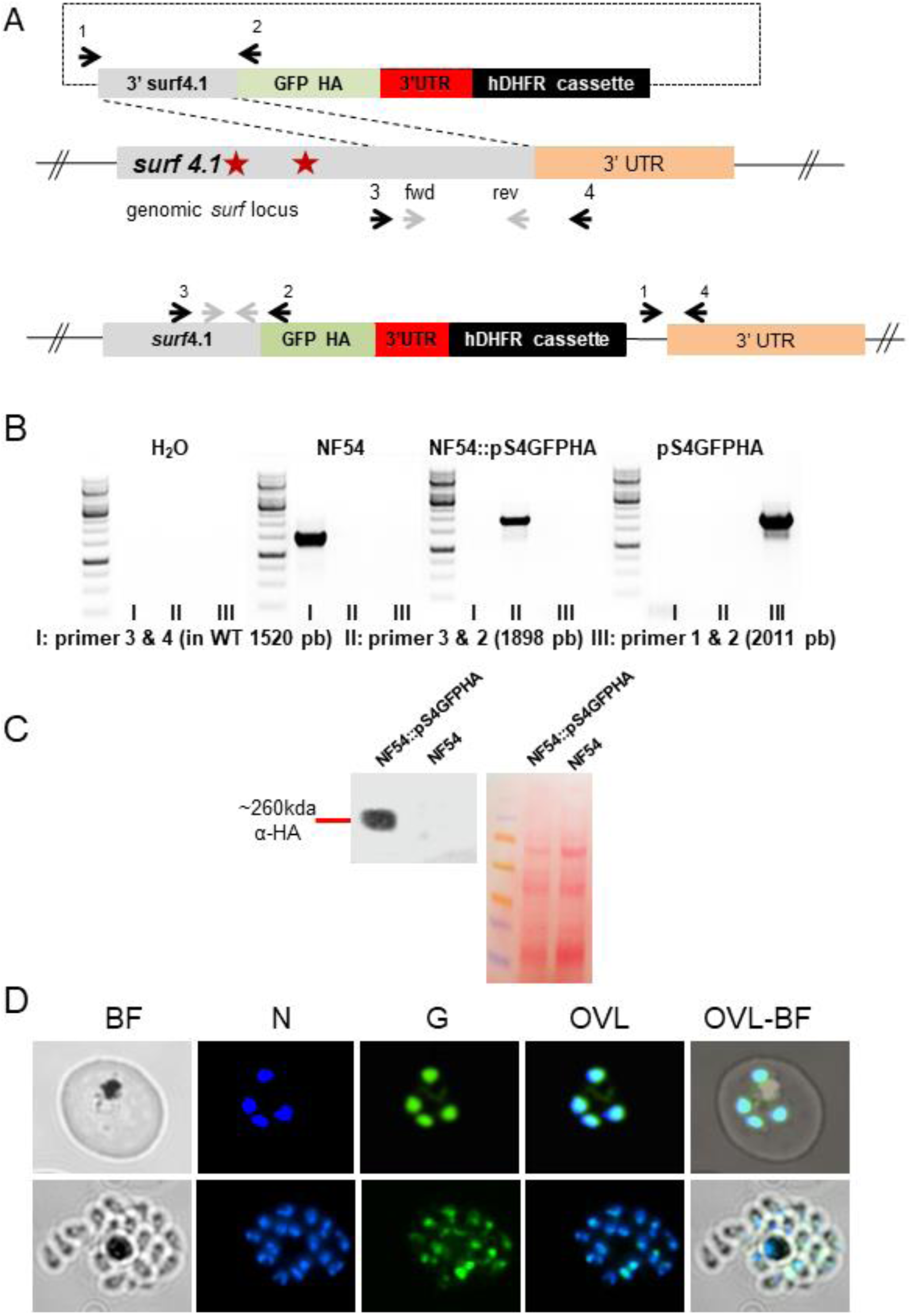
Tagging of SURFIN 4.1 with GFP and HA leads to green fluorescent parasites. In **A**, the proposed model for single crossover recombination of the plasmid pS4-GFP-HA is shown. The arrows indicate the localization of oligonucleotides used for PCR. Grey arrows indicate the oligonucleotides which were used to create the knockin-homology region in pS4-GFP-HA. In **B,** PCR with the primers indicated above show knockin of the construct into the *surf*4.1 locus. The primer combinations used in PCRs and the amplicon sizes are indicated. Note that primer pair II results in an amplicon solely in knocked-in parasites. In **C**, Western blot of NF54::pS4-GFP-HA and NF54 show the full-length, tagged SURFIN4.1 in schizont stage in the transfectant line but not NF54 wildtype parasites. On the right, the loading of each lane is demonstrated by Ponceau-staining. In **D**, Fluorescence microscopy with a schizont from the parasite line NF54::pS4GFPHA I, bright field, II, nuclear staining using DAPI, III, GFP-tagged SURFIN4.1 and IV, overlay of I to III.

### A novel stop codon abrogates production of SURFIN4.1-GFP-HA

To examine if the presence of an artificially introduced stop codon (TAA) could be introduced in the *surf*4.1 ORF, additional transfectant lines were prepared (Figure 2). The first one was transfected with the plasmid pS4(TAA)GFPHAglms-TK and the second with pS4GFPHAglmS-TK. Both plasmids differ in i) one single base pair which introduces a stop codon in the middle of the first homology region which contains the 3’-ORF of *surf*4.1 (Figure 2 and Supplementary Figure 3) and ii) the 3’-ORF homology region of pS4GFPHAglmS-TK is shorter. Following outgrowth of the parasite lines and one cycling without/with WR99210 as described above, the resulting parasite lines were treated with ganciclovir. Integration into the genome would then insert a novel stop codon in the predicted translated ORF. However, after ganciclovir treatment no parasites were recovered from the pS4(TAA)GFPHAglms-TK transfected line, suggesting that the insertion of a new stop codon into the genomic sequence of *surf*4.1 severely interfered with the survival of the parasite during blood stage. In contrast, the transfectant line bearing the plasmid pS4GFPHAglmS-TK was readily recovered and showed rapid integration into the *surf*4.1 locus (Figure 2), as shown by PCR with integration-specific oligonucleotides. Of note, while the stop-codon containing parasite line failed to show a knockin genotype (Figure 2, seen as an amplicon with oligo pair III, gels with amplicons from genomic DNA of NF54::pS4(TAA)GFPHAglmS-TK+Gcv, with and without WR), the specific presence of a knockin genotype with the absence of an amplicon for the TK cassette indicated a successful knockin resulting in the NF54::pS4GFPHAglmS-TK line. As controls, genomic DNA from NF54 and the transfection plasmid pS4GFPHAglmS-TK resulted solely in the expected PCR products for the presence of the 1303 bp amplicon (oligos 3 and 4) or the TK cassette (931 bp, oligos 1 and 2). Southern blot analysis using BamH1-digested genomic DNA from the ganciclovir-treated NF54::pS4GFPHAglmS-TK line confirmed the successful knockin and the expected 7786 bp digestion product, stemming from a BamH1 site upstream of the knockin fragment of *surf*4.1 and a second site in the hDHFRcassette (Figure 2 C). In contrast, the transfection plasmid was visualized in its linearized form, detected as a 11380 bp fragment.

**Figure 2:**
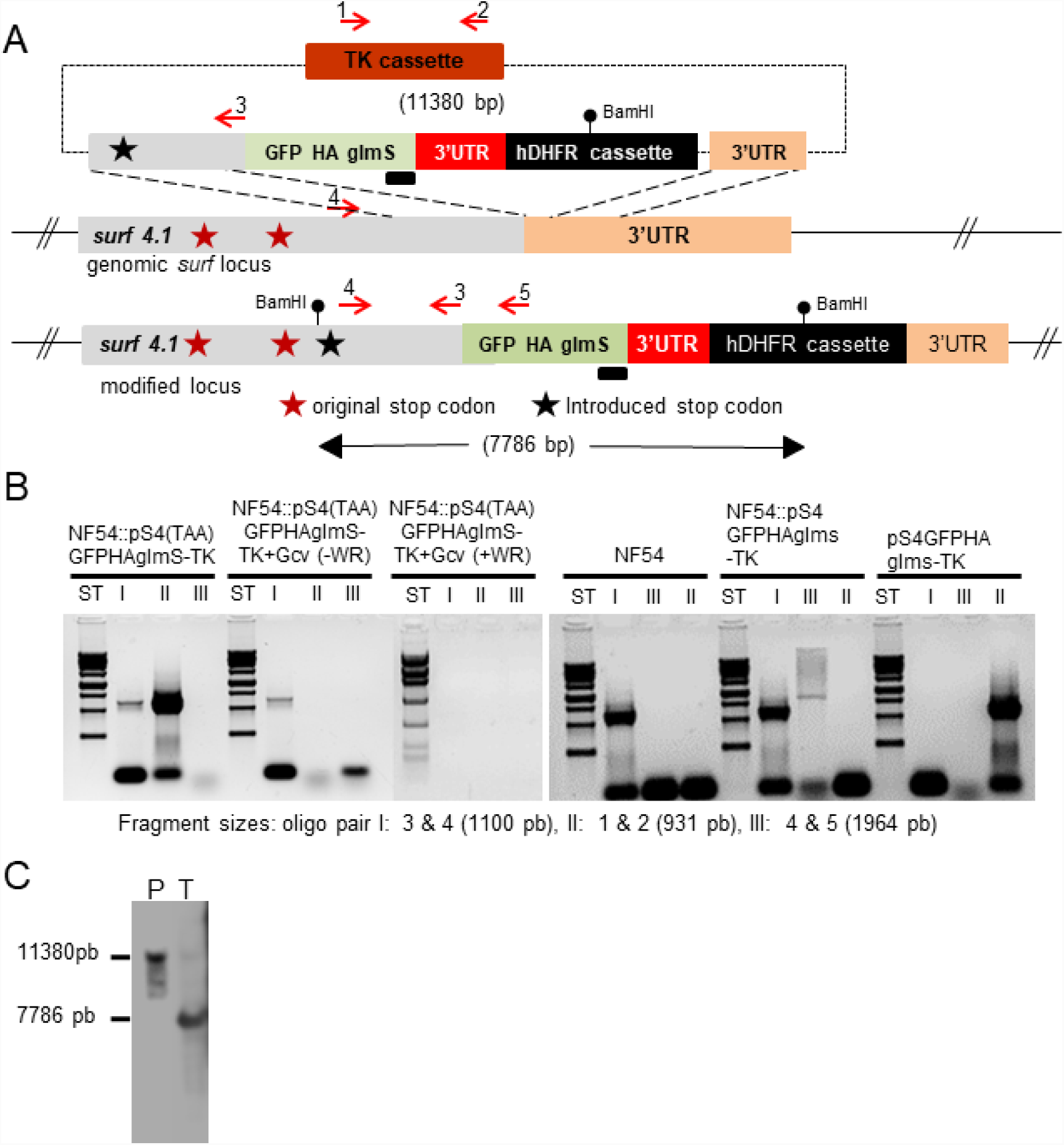
A construct containing a novel stop codon in the 3’ part of the *surf*4.1 ORF is refractory to knockin. In **A**, the proposed model for double crossover recombination of the plasmid pS4(TAA)-GFPHAglmS-TK is shown. The arrows indicate the localization of oligonucleotides used for PCR and each red star indicates stop codons predicted in PlasmoDB and a black star indicates the position of the novel introduced stop codon. In **B,** genomic DNA from transfectant lineages treated or not with Ganciclovir (Gcv) is characterized by PCR by the indicated primer combinations. Note that primer pair I amplifies fragments from wild type and knocked-in constructs. Amplification products using material from the parasite line transfected with pS4(TAA)-GFPHAglmS-TK and treated with Gcv and afterwards with WR for 4 days are shown in the 3rd panel from the left and appear devoid of any amplifiable material. The panels on the right show amplicons from either untransfected NF54 genomic DNA, or transfectants using the pS4-GFPHAglmS-TK before and after Gcv treatment. In **C**, Southern blot analysis using the glmS specific probe (black bar in **A**) shows integration as expected of the constructs and the virtual absence of episomal material. “P” contains BamHI-linearized plasmid and “T” contains 10 µg BamHI restricted genomic DNA from NF54::pS4-GFPHAglmS-TK.

To further confirm if *surf*4.1 is essential, a conventional knockout by gene disruption was attempted, using transfecting plasmid pS4KO (Supplementary Figure 3). Again, the transfected parasite line did not survive ganciclovir treatment even after three drug-on/off cycles. It was also impossible to detect any signal for integration in *surf*4.1 by PCR on genomic DNA of the transfected parasite line (data not shown).

### glmS-Glucosamine-mediated knockdown of *surf*4.1 strongly inhibits schizont development

Given its probable importance for the survival of parasites during blood stages, we tested if a transcript knockdown led to a measurable growth phenotype. In previous studies, the SURFIN4.1 protein knockdown to approximately 20% of the normal SURFIN4.1 level resulted only in an increase of the *surf*4.1 transcript, but no apparent growth phenotype (Macedo-Silva et al., 2017). To test for a growth phenotype, the integrated parasite line used as a control line in the previous experiment (Figure 3) was submitted to glucosamine-6-phosphate treatment. First, the efficiency of knockdown was monitored and RT-qPCR showed an approximately 10fold decrease in the *surf*4.1 message (Figure 3) while other surf transcripts were either silenced or not altered (Supplementary Figure 4). Glucosamine treatment led also to a decrease in the quantity of the SURFIN4.1-GFP-HA protein, while the PTEX polypeptide (loading control) was not strongly influenced. Image analysis indicated a 70% knockdown of SURFIN4.1-GFP-HA after 24 h treatment with glucosamine (Figure 3 C). The viability of parasites during knockdown was also tested. During a 96 h treatment with glucosamine-6-phosphate, a significant growth inhibition started apparently at the time point of schizont lysis and reinvasion at around 40 h post treatment initiation (Figure 3 D), while treatment of untransfected NF54 parasites only slightly suppressed parasite multiplication (Figure 3 E).

**Figure 3:**
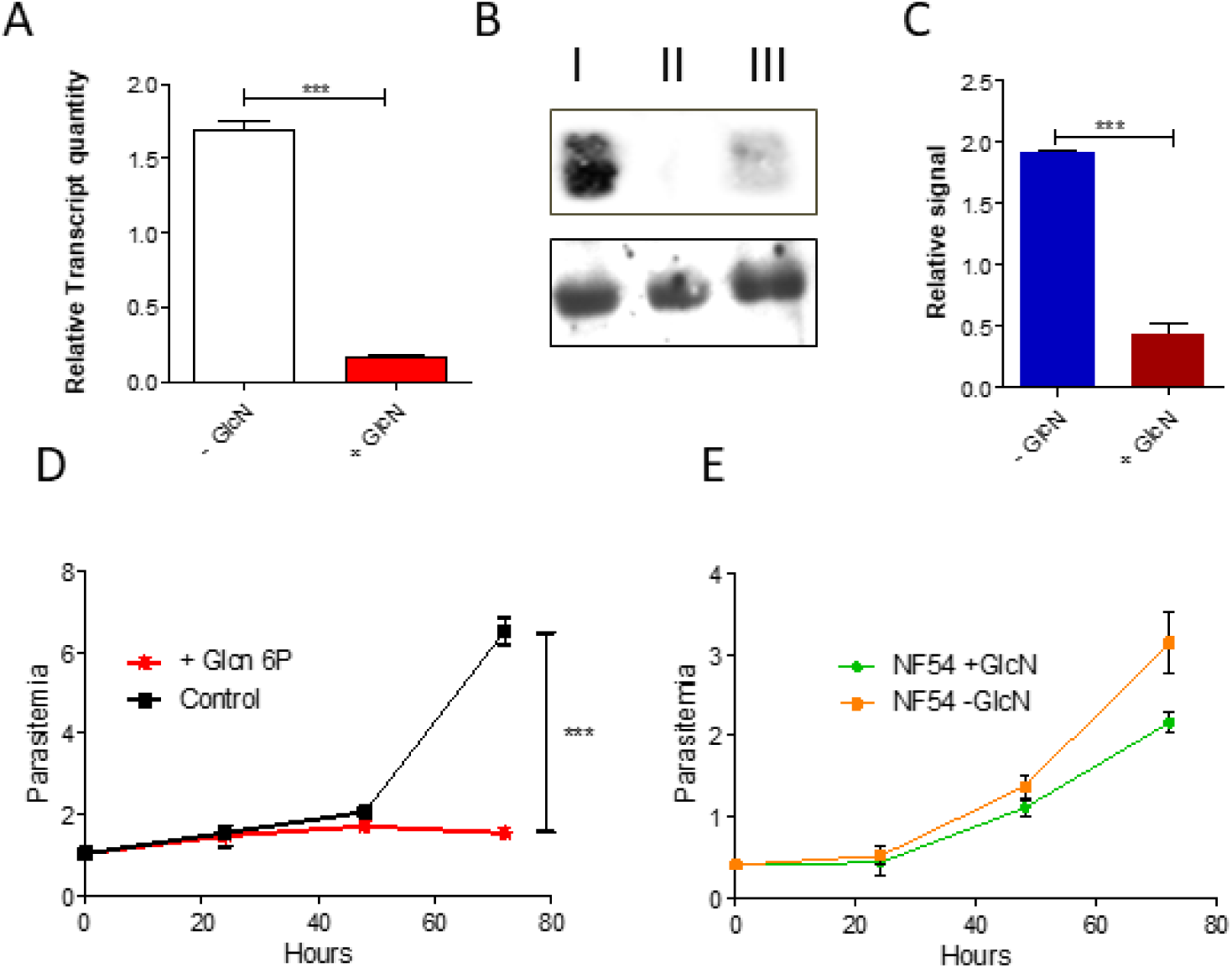
Transcript knockdown of *surf*4.1 results in a decrease of *surf*4.1 transcripts, SURFIN4.1-GFP-HA protein and impairs parasite growth. In **A**, knockdown of the *surf*4.1 transcript measured by RT-qPCR in NF54::pS4-GFPHAglmS-TK transfectants treated or not with 2.2 mM glucosamine-phosphate in relation to the endogenous control seryl-tRNA ligase. In **B-** Western blot of NF54::pS4-GFPHAglms and NF54 in schizont stage and in the presence and absence of glucosamine-phosphate, showing SURFIN4.1-GFP-HA (representative of 2 individual experiments) and untransfected NF54 (no detectable signal in 260 kDa, due to the absence of the HA tag). The loading control below was done with anti-PTEX150. In **C**, densitometry analysis of the observed signals in B using ImageJ, normalized against the PTEX150 signal. Three asterisks mean highly significant differences (*p*= 0.0006, Student’s T-test, with a mean of the difference of 1.472 relative units). **D:** Effect on the parasitemia of GlcN and subsequent absence of SURFIN4.1is shown. Error bars indicate the standard deviation of transcription of each surf gene in individual triplicates. Three asterisks mean highly significant differences (p < 0.0001, Two-Way ANOVA). **E**: A slight negative influence of Glucosamine phosphate on the growth of wildtype NF54 cultures is shown.

### Disruption of *surf*4.1 impairs schizogony and parasite proliferation

We then monitored the morphologic effects of parasites submitted to glmS-mediated SURFIN4.1 knockdown. For this, SURFIN4.1-GFP-HA-expressing parasites were cultivated in the presence of 2.2 mM of glucosamine starting in ring stage (6 hpi). A stalling of parasite growth was observed (Figure 4) and the knocked down parasite seems to be unable to perform a proper merozoite development, especially in late trophozoites/schizonts. Longer treatment indicated that the stalled parasites did not complete schizogony and thus were unable to reinvade. When parasite forms were counted, strong and significant inhibition of the schizont development was observed after knockdown of the *surf*4.1 transcript. Longer cultivation in the presence of glucosamine-6-phosphate was impossible and parasites disappeared while untreated parasites proliferated normally (data not shown).

**Figure 4:**
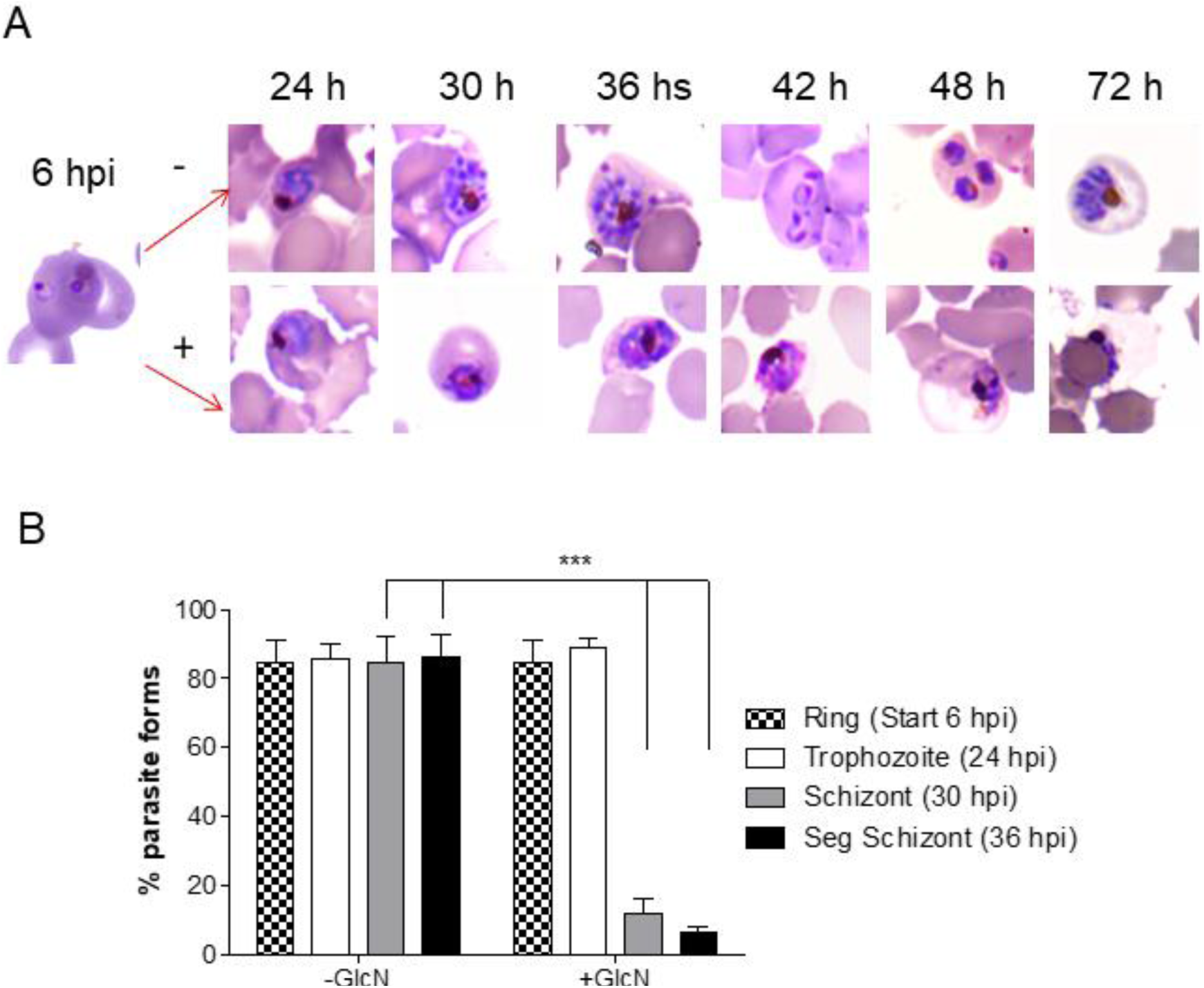
Parasites with knocked-down SURFIN4.1 show a strongly reduced capacity of parasites to progress to mature schizonts. **A**: Highly synchronized parasites (2 sequential cycles of floating/sorbitol treatment) were submitted to Glucosamine phosphate treatment or not. The progression of intraerythrocytic parasite development was examined by thin blood smears stained with Giemsa (representative data of 3 experiments). **B**: Counting of slides resulted in the observation that parasites stop to develop in late-stage trophozoites.

### SURFIN4.1 probably interacts with tubulin and hsp90

Quintana et al. detected that SURFIN4.2 from the *P. falciparum* IT strain forms a complex between RON4 and GLURP and are probably involved in the formation of the parasite vacuole at reinvasion (Quintana et al., 2018). If SURFIN4.2 from the IT/FCR3 strain and SURFIN4.1 of the NF54 line are functionally equivalent, this interaction may be confirmed by mass spectrometry analysis. Using antiGFP-sepharose, co-immunoprecipitating proteins were trypsin-digested and analyzed by OrbiTrap mass spectrometry. As a result, a number of proteins were detected, including tubulin and heat shock protein 70 and 90 (Figure 5), but not RON4 or GLURP. To support a possible interaction between β-tubulin and SURFIN4.1, immunofluorescence microscopy was employed and a partial overlay of the fluorescence of SURFIN4.1-GFP-HA and β-tubulin was visualized (Figure 5). Interestingly, the knockdown led to the absence not only of GFP but also β-tubulin fluorescence. Also, in parasites under knockdown conditions, a decrease in the production of β-tubulin is notable compared to parasites without glucosamine 6 phosphate treatment (Figure 5 D). In the same samples, the quantity of histone 3 is not influenced. Beta-tubulin is an essential part of microtubules which, in turn, form an integral part of the cytoskeleton that is a dynamic set of long and thin fibers that contribute to the transport of intracellular components and in cell division. The lack of SURFIN4.1 seems to lead to a breakdown of the merozoite-forming process and a concomitant decrease of DAPI fluorescence. This may point to a lack of mitotic activity. In order to establish an influence of glucosamine treatment and SURFIN4.1 knockdown on the replication process, genomic DNA of an identical number of glucosamine-treated and untreated parasites was quantified. As shown in Figure 6, 20 h after the onset of glucosamine treatment (starting in ring stage parasites), an increase of the DNA quantity could be identified compared to the initial sample. However, the DNA quantities in glucosamine-treated or untreated parasites were not significantly different, pointing to the view that replication itself may not have been influenced. Taken together, the lack of SURFIN4.1 severely interferes with schizont development and this may occur by influencing intracellular trafficking.

**Figure 5:**
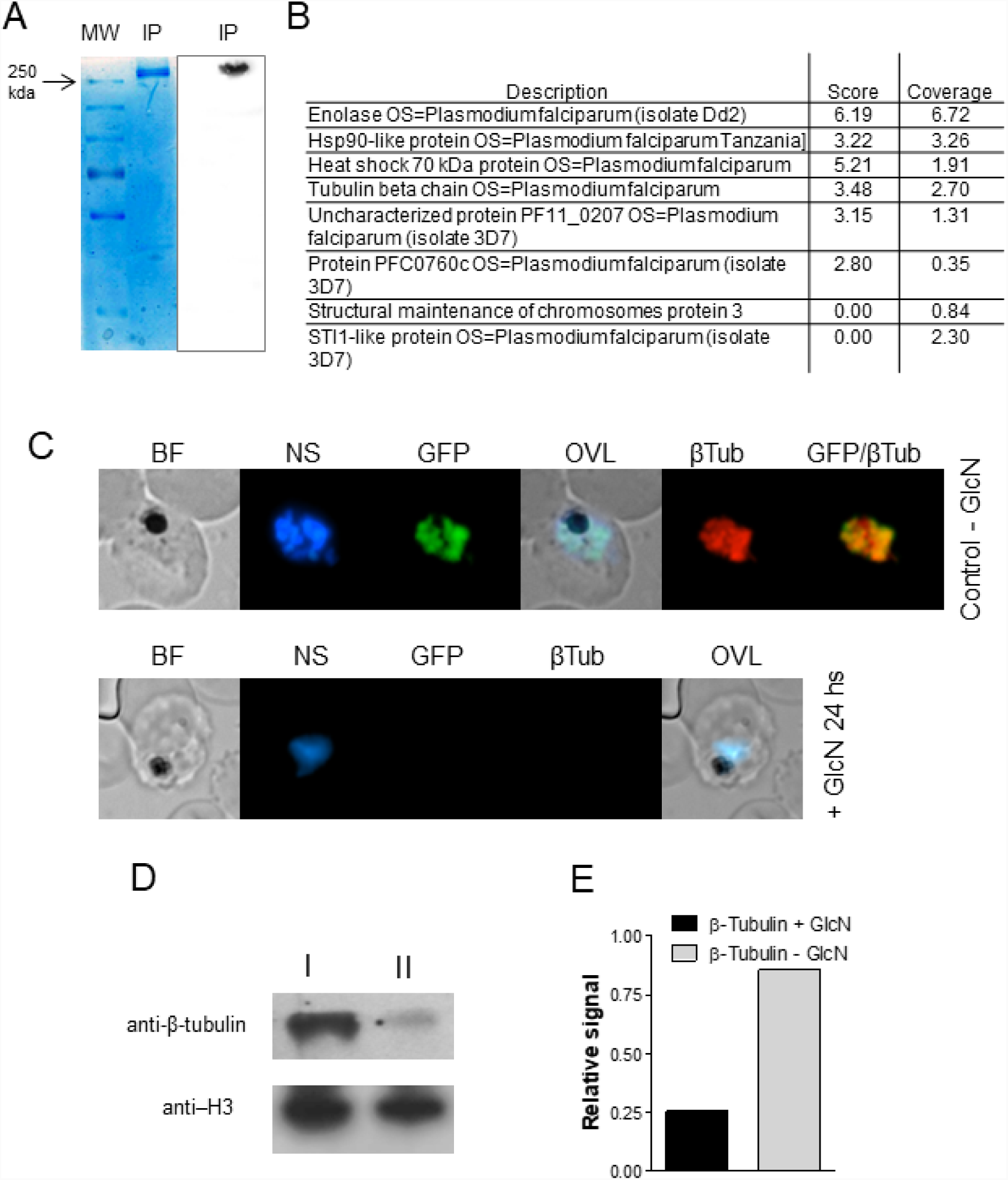
Co-IP by Pull-down and mass spectrometry of SURFIN4.1 and Immunofluorescence showing the effect on the tubulin structure and nuclear increase and division (DAPI mark) after disruption of SURFIN4.1. In A, left side, Colloidal Coomassie blue stained SDS-PAGE (8%) was loaded with a third of the eluted material from immunoprecipitated lysed parasites (2 ml compacted red blood cells at 8% parasitemia late trophozoites/schizonts) and a second third was run on the same gel, transferred onto nitrocellulose and later GFP-containing proteins were detected with antiGFP as described. The third part of the eluted material was trypsin digested and analyzed by mass spectrometry resulting in the co-precipitating protein species as shown. Listed are only proteins which produced peptide fragments coincident with unique peptides from the *P. falciparum* databank (**B**). The complete curated output from ProteinDiscovery can be accessed in Supplementary Table 2. In **C**, parasites were examined for SURFIN4.1-GFP and beta-tubulin colocalization. Note that treatment with glucosamine phosphate (GlcN) strongly decreases both SURFIN4.1-GFP and beta-tubulin detection. In **D**, western blot detection with the indicated antibodies revealed a strong decrease in beta-tubulin presence while Histone 3 is not influenced. Quantification using ImageJ (**E**) resulted in only around 30% of the beta-tubulin signal in GlcN treated parasites.

**Figure 6:**
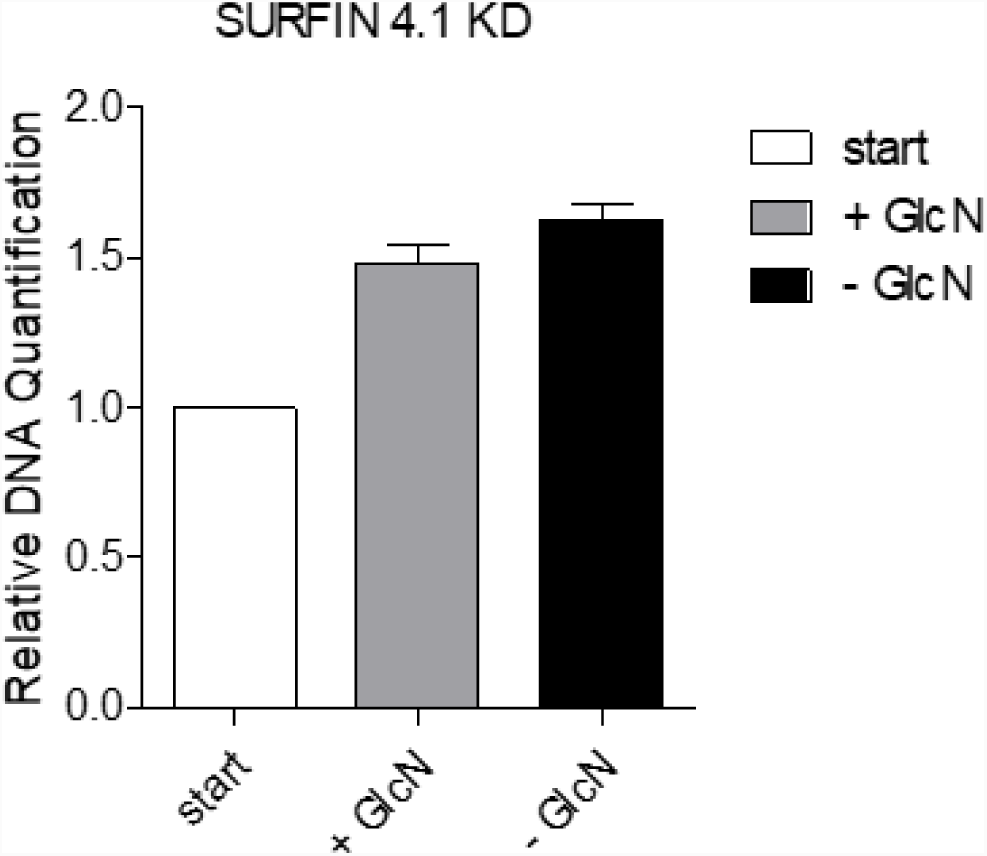
Knockdown of SURFIN4.1-GFP does not interfere with DNA replication. Parasites were treated or not with 2.5 mM GlcN for 24 h starting in early ring stage. An equal number of parasites had their genomic DNA extracted at timepoint 0 and 24 h. Ct values were obtained for the single copy gene (seryl-tRNA ligase) and compared between each other, assuming that each Ct unit difference equals the double amount of input DNA. Experiments were done in triplicate.

## Discussion

The *surf* gene family and their orthologs are present in many primate species of *Plasmodium*. Specifically, *surf*4.1 is localized in a highly syntenic region in different *Plasmodium falciparum* strains, but also in *Plasmodium gaboni* and *Plasmodium reichenowi,* where it is localized between the genes encoding PHISTb and Reticulocyte Binding protein 1 homolog. Intriguingly, and despite its apparent conservation, *surf*4.1 is annotated in several sequenced strains as a pseudogene, showing stop codons along its predicting open reading frame. Also, slightly different patterns of splicing sites were predicted: While there is apparently no splicing in the *surf*4.1 variant of the *P. falciparum* GA01 strain, two introns were predicted for the SN01 strain (PlasmoDB 41). Another strain (ML01 strain), also with no predicted intron, shows *surf*4.2 instead of *surf*4.1 in the syntenic region. Here, we confirmed that the stop codons were also present in the transcripts of *surf*4.1. Together with the observation that in the 3D7 strain there are in total 160 genes annotated as pseudogenes of which for 29 exist proteomic evidence (including SURFIN4.1), we conclude that readthrough of stop codons is a frequent event. However, which amino acid is incorporated at the place where a stop codon occurs is unknown and was not further addressed. Intriguingly, there must be an adaptation to the read-through of stop codons, since an artificially introduced stop codon resulted in a non-viable phenotype. A more detailed analysis, perhaps including a three-dimensional structure of the transcript, may reveal patterns which allow the ribosome to read through stop codons. In many organisms, the sequence surrounding stop codons clearly plays a role in the decision if a read through occurs or not (Dabrowski et al., 2015). In several organisms, stop codons are translated into seleno-cysteines (reviewed in (Rodnina et al., 2017)). In *P. falciparum*, a selenocysteine tRNA which recognizes the UGA stop codon was described (Mourier et al., 2005). Later, Lobanov and colleagues predicted four seleno-proteins, none of which is SURFIN4.1 (Lobanov et al., 2006). Regarding *surf*4.1 splicing variants, other authors also already documented uncommon splicing variants of this gene (Zhu et al., 2013), and one resulted in the deletion of the tryptophan-rich domains at the C-terminus of the 3D7 line. These authors also tested different domains for their contribution to the subcellular trafficking of SURFIN4.1. However, in none of their constructs, the complete, stop-codon containing open reading frame of *surfin*4.1 was tested, as was successfully done in this work. Interestingly, Zhu and colleagues found a truncated variant of SURFIN4.1 using an antibody raised against the N-terminal domain, but no full-length protein was shown in their analyses (Zhu et al., 2013) using the untransfected 3D7A line. However, the corresponding blot appears not to show very large proteins. In the same study, SURFIN4.1 was detected in different sites depending on which domains were included for transgene expression in *Plasmodium*, while the herein used GFP-tagged variant localized clearly and exclusively to merozoites. The reason for this discrepancy is not clear and perhaps strain differences are responsible for the observed differences. It is probable that we missed truncated variants in our experiments since only full-length proteins were detectable in the experimental setup. In the same way, it is possible that both the full length and truncated versions coexist and play different biological roles in the parasite dependent on their localization, which may be determined by the presence or absence of the C-terminal domain of the protein.

Regarding the function of the antigen SURFIN4.1, we have shown in a previous study that full-length SURFIN4.1 could be knocked down without triggering any growth phenotype (Macedo-Silva et al., 2017) and the only effect observed was the increase of its transcript. Initially, and to further explore this phenomenon, we created a parasite line where the transcript could be knocked down. Surprisingly, the knockdown of the *surf*4.1 transcript had profound effects on the schizont development and the observed phenotypes showed a stalling of the parasite in the late trophozoite/early schizont stage, which was not observed in the SURFIN4.1-protein knockdown. When comparing the knockdown efficiency of the full-length protein using either the degron (destabilizing domain variant “24” (de Azevedo et al., 2012; Macedo-Silva et al., 2017)) or the transcript knockdown (Prommana et al., 2013) applied here, the overall knockdown efficiency observed in western blots was comparable. One explanation may be that truncated forms as observed by Zhu and colleagues have an important role in the effects observed here, while the full-length protein has perhaps only a partial and accessory effect. The transcript knockdown would suppress all resulting forms of the SURF4.1 protein, while the degron-mediated knockdown only interferes with the full-length protein, which possesses the destabilizing domain while truncated forms do not. The use of an antibody against an N-terminal region – not available in this study - of SURFIN4.1 may elucidate this question.

We then addressed the function of SURFIN4.1. In a previous study, Quintana and colleagues provided evidence that SURFIN4.2 interacts with GLURP and RON4, and blockage of SURFIN by antibodies against its ectodomain led to a small but measurable decrease in the reinvasion of merozoites into erythrocytes (Quintana et al., 2018). No data, however, are available what missing SURFIN4.1 or IT/FCR SURFIN4.2 would cause in the formation of the postulated complex. By knocking down the *surf*4.1 transcript, a profound perturbation of merozoite formation and stalling of schizont development was observed, reminiscent of a block in cell cycle progression. However, and documented in Figure 6, no significant interference in DNA replication was found in ∼24 h post reinvasion parasites treated or not with glucosamine-phosphate, indicating that the observed stalling in early schizont stage was possibly due to the structural or metabolic hindrance of merozoite formation, and not interference in genome replication. In a recent study, genome-wide integration of insertions using the piggy-bac approach was applied to characterize genes which were possibly essential in *P. falciparum* (Zhang et al., 2018). In this approach, *surf*4.1 was deemed amenable to insertion without hampering the growth of the parasite. This apparent conflict may be explained by the fact that piggyBac-mediated insertion occurred in sequences at 400 nt near the 3’ end of the *surf*4.1 reading frame and therefore after the stop codons which lead to the possibly essential truncated forms of SURFIN4.1, and also after the inserted stop codon introduced in pS4(TAA)GFPHAglmS-TK. No other successful insertion was observed in the remaining approximately 6000 base pairs upstream ((Zhang et al., 2018), compare supplementary data, Table S1). RNAseq results deposited by different authors may give a hint at what stage different SURFINs possibly exert functions. In the dataset published by Otto and colleagues (Otto et al., 2010) the highest transcript quantities in trophozoites and schizonts were also found for *surf*4.1, while another *surf* gene (PF3D7_0113100) was strongly detected in ookinetes. In another study which approached transcript bound to polysomes, *surf*4.1 was also the most abundantly present surf transcript in schizonts (Bunnik et al., 2013). In yet another experiment monitoring transcripts in the 3D7 strain, *surf*4.1 was also the most abundantly found transcript in trophozoites and schizonts (López-Barragán et al., 2011). Collectively, this reinforces a role for SURFIN4.1 at least in the 3D7/NF54 parasite lines.

In order to detect interacting proteins, mass spectrometry was used to identify proteins which interact with GFP-tagged SURFIN4.1. While heat shock proteins 70, 90 and beta-tubulin were detected, neither RON4 nor GLURP could be identified. This may be due to the fact that the complex between these factors is built only in later stages which are not formed when SURFIN4.1 is knocked down, or else, full-sized SURFIN4.1 does not interact with these proteins, differently from IT/FCR3’s SURFIN4.2. We are currently addressing the question of why enolase is possibly interacting with SURFIN4.1. In other studies, plasmodial enolase was identified as a multifunctional protein, found in different subcellular localizations (Pal Bhowmick et al., 2009) and possibly interacting with a number of different factors, apparently including SURFIN4.1. When we tried to confirm the colocalization of SURFIN4.1 and beta-tubulin, knocked-down parasites did not reveal any signal of beta-tubulin at all, while a partially overlapping fluorescence signal was observed for schizont stage parasites kept in the absence of glucosamine phosphate. In immunoblots, beta-tubulin also appeared in decreased amounts, indicating that less SURFIN4.1 also somehow leads to the suppression of beta-tubulin production. Alternatively, knocked down parasites are unable to reach the phase when beta-tubulin is strongly expressed. Of note, the expression of beta-tubulin varies during the blood stage parasite development and beta-tubulin transcription is strongest in stages older than 20 h post reinvasion (Otto et al., 2010) and the protein is weakly visible in younger forms (Fennell et al., 2008). In two hybrid-studies, SURFIN4.1 appeared to interact predominantly with DNA binding proteins such as zinc-finger nucleases, DNA binding chaperones, SET10 – a histone lysine N-methyl transferase, putative kinetochore-suppressor protein, but also a Maurer’s cleft protein (ETRAMP) and a putative ABC transporter, besides others (LaCount et al., 2005). These may not have been detected in our proteomic analyses due to their low abundance.

Taken together, while the exact function of SURFIN4.1 remains elusive we have shown that this unusual protein possesses essential functions during the early schizogony phase of intraerythrocytic development. Taking in account the differential expression of *surf* alleles in different stages of *P. falciparum* forms including sporozoites and ookinetes, the functions of each *surf* allele area probably non-redundant unlike other multigene families such as *var* and *rif* and may reveal targets of intervention against this still devastating disease.

## Supporting information

This file contains 4 supplementary Figures.

This file contains two Supplementary Tables.

## Acknowledgments

This work was supported by FAPESP grants 2015/17174-7 and 2017/24267-7. TMS and RBDA were supported by CNPq fellowships. GW is a CNPq research fellow. The proteomic analysis was performed at the CEFAP at ICB-USP.

## Authors contributions

TMS and RBDA performed experiments. GW and TMS conceived the experimental outline and wrote the manuscript.

## Literature

de Azevedo, M.F., Gilson, P.R., Gabriel, H.B., Simões, R.F., Angrisano, F., Baum, J., Crabb, B.S., and Wunderlich, G. (2012). Systematic Analysis of FKBP Inducible Degradation Domain Tagging Strategies for the Human Malaria Parasite Plasmodium falciparum. PLoS One 7, e40981.

Baruch, D.I., Pasloske, B.L., Singh, H.B., Bi, X., Ma, X.C., Feldman, M., Taraschi, T.F., and Howard, R.J. (1995). Cloning the P. falciparum gene encoding PfEMP1, a malarial variant antigen and adherence receptor on the surface of parasitized human erythrocytes. Cell 82, 77–87.

Bunnik, E.M., Chung, D.-W., Hamilton, M., Ponts, N., Saraf, A., Prudhomme, J., Florens, L., and Le Roch, K.G. (2013). Polysome profiling reveals translational control of gene expression in the human malaria parasite Plasmodium falciparum. Genome Biol. 14, R128.

Cheng, Q., Cloonan, N., Fischer, K., Thompson, J., Waine, G., Lanzer, M., and Saul, A. (1998). stevor and rif are Plasmodium falciparum multicopy gene families which potentially encode variant antigens. Mol Biochem Parasitol 97, 161–176.

Dabrowski, M., Bukowy-Bieryllo, Z., and Zietkiewicz, E. (2015). Translational readthrough potential of natural termination codons in eucaryotes--The impact of RNA sequence. RNA Biol. 12, 950–958.

Duraisingh, M.T., Triglia, T., and Cowman, A.F. (2002). Negative selection of Plasmodium falciparum reveals targeted gene deletion by double crossover recombination. Int. J. Parasitol. 32, 81–89.

Fennell, B.J., Al-shatr, Z.A., and Bell, A. (2008). Isotype expression, post-translational modification and stage-dependent production of tubulins in erythrocytic Plasmodium falciparum. Int. J. Parasitol. 38, 527–539.

Fernandez, V., Hommel, M., Chen, Q., Hagblom, P., and Wahlgren, M. (1999). Small, clonally variant antigens expressed on the surface of the Plasmodium falciparum-infected erythrocyte are encoded by the rif gene family and are the target of human immune responses. J. Exp. Med. 190, 1393–1404.

Freitas-Junior, L.H., Bottius, E., Pirrit, L.A., Deitsch, K.W., Scheidig, C., Guinet, F., Nehrbass, U., Wellems, T.E., and Scherf, A. (2000). Frequent ectopic recombination of virulence factor genes in telomeric chromosome clusters of P. falciparum. Nature 407, 1018–1022.

Gitaka, J.N., Takeda, M., Kimura, M., Idris, Z.M., Chan, C.W., Kongere, J., Yahata, K., Muregi, F.W., Ichinose, Y., Kaneko, A., et al. (2017). Selections, frameshift mutations, and copy number variation detected on the surf 4.1 gene in the western Kenyan Plasmodium falciparum population. Malar. J. 16, 98.

Gölnitz, U., Albrecht, L., and Wunderlich, G. (2008). Var transcription profiling of Plasmodium falciparum 3D7: assignment of cytoadherent phenotypes to dominant transcripts. Malar J 7, 14.

Hasenkamp, S., Russell, K.T., and Horrocks, P. (2012). Comparison of the absolute and relative efficiencies of electroporation-based transfection protocols for Plasmodium falciparum. Malar. J. 11, 210.

van der Heyde, H.C., Nolan, J., Combes, V., Gramaglia, I., and Grau, G.E. (2006). A unified hypothesis for the genesis of cerebral malaria: sequestration, inflammation and hemostasis leading to microcirculatory dysfunction. Trends Parasitol 22, 503–508.

Hiller, N.L., Bhattacharjee, S., van Ooij, C., Liolios, K., Harrison, T., Lopez-Estraño, C., and Haldar, K. (2004). A host-targeting signal in virulence proteins reveals a secretome in malarial infection. Science (80-.). 306, 1934–1937.

LaCount, D.J., Vignali, M., Chettier, R., Phansalkar, A., Bell, R., Hesselberth, J.R., Schoenfeld, L.W., Ota, I., Sahasrabudhe, S., Kurschner, C., et al. (2005). A protein interaction network of the malaria parasite Plasmodium falciparum. Nature 438, 103–107.

Lambros, C., and Vanderberg, J.P. (1979). Synchronization of Plasmodium falciparum erythrocytic stages in culture. J Parasitol 65, 418–420.

Larkin, M.A., Blackshields, G., Brown, N.P., Chenna, R., McGettigan, P.A., McWilliam, H., Valentin, F., Wallace, I.M., Wilm, A., Lopez, R., et al. (2007). Clustal W and Clustal X version 2.0. Bioinformatics 23, 2947–2948.

Lelievre, J., Berry, A., and Benoit-Vical, F. (2005). An alternative method for Plasmodium culture synchronization. Exp Parasitol 109, 195–197.

Ljungström, I., Perlmann, H., Schlichtherle, M., Scherf, A., and Wahlgren, M. (2008). Methods in Malaria Research (Manassas, VA: MR4/ATCC, BioMalPar).

Lobanov, A. V, Delgado, C., Rahlfs, S., Novoselov, S. V, Kryukov, G. V, Gromer, S., Hatfield, D.L., Becker, K., and Gladyshev, V.N. (2006). The Plasmodium selenoproteome. Nucleic Acids Res. 34, 496–505.

López-Barragán, M.J., Lemieux, J., Quiñones, M., Williamson, K.C., Molina-Cruz, A., Cui, K., Barillas-Mury, C., Zhao, K., and Su, X. (2011). Directional gene expression and antisense transcripts in sexual and asexual stages of Plasmodium falciparum. BMC Genomics 12, 587.

Macedo-Silva, T., Araujo, R.B.D., Meissner, K.A., Fotoran, W.L., Medeiros, M.M., de Azevedo, M.F., and Wunderlich, G. (2017). Knockdown of the Plasmodium falciparum SURFIN4.1 antigen leads to an increase of its cognate transcript. PLoS One 12, e0183129.

Maier, A.G., Rug, M., O’Neill, M.T., Brown, M., Chakravorty, S., Szestak, T., Chesson, J., Wu, Y., Hughes, K., Coppel, R.L., et al. (2008). Exported proteins required for virulence and rigidity of Plasmodium falciparum-infected human erythrocytes. Cell 134, 48–61.

Marti, M., Good, R.T., Rug, M., Knuepfer, E., and Cowman, A.F. (2004). Targeting malaria virulence and remodeling proteins to the host erythrocyte. Science (80-.). 306, 1930–1933.

Milner, D.A. (2018). Malaria Pathogenesis. Cold Spring Harb. Perspect. Med. 8, a025569.

Mourier, T., Pain, A., Barrell, B., and Griffiths-Jones, S. (2005). A selenocysteine tRNA and SECIS element in Plasmodium falciparum. RNA 11, 119–122.

Mphande, F.A., Ribacke, U., Kaneko, O., Kironde, F., Winter, G., and Wahlgren, M. (2008). SURFIN4.1, a schizont-merozoite associated protein in the SURFIN family of Plasmodium falciparum. Malar. J. 7, 116.

Ochola, L.I., Tetteh, K.K.A., Stewart, L.B., Riitho, V., Marsh, K., and Conway, D.J. (2010). Allele Frequency-Based and Polymorphism-Versus-Divergence Indices of Balancing Selection in a New Filtered Set of Polymorphic Genes in Plasmodium falciparum. Mol. Biol. Evol. 27, 2344–2351.

Otto, T.D., Wilinski, D., Assefa, S., Keane, T.M., Sarry, L.R., Böhme, U., Lemieux, J., Barrell, B., Pain, A., Berriman, M., et al. (2010). New insights into the blood-stage transcriptome of *Plasmodium falciparum* using RNA-Seq. Mol. Microbiol. 76, 12–24.

Pal Bhowmick, I., Kumar, N., Sharma, S., Coppens, I., and Jarori, G.K. (2009). Plasmodium falciparum enolase: stage-specific expression and sub-cellular localization. Malar. J. 8, 179.

del Portillo, H.A., Fernandez-Becerra, C., Bowman, S., Oliver, K., Preuss, M., Sanchez, C.P., Schneider, N.K., Villalobos, J.M., Rajandream, M.A., Harris, D., et al. (2001). A superfamily of variant genes encoded in the subtelomeric region of Plasmodium vivax. Nature 410, 839–842.

Prommana, P., Uthaipibull, C., Wongsombat, C., Kamchonwongpaisan, S., Yuthavong, Y., Knuepfer, E., Holder, A.A., and Shaw, P.J. (2013). Inducible Knockdown of Plasmodium Gene Expression Using the glmS Ribozyme. PLoS One 8, e73783.

Quintana, M.D.P., Ch’ng, J.-H., Zandian, A., Imam, M., Hultenby, K., Theisen, M., Nilsson, P., Qundos, U., Moll, K., Chan, S., et al. (2018). SURGE complex of Plasmodium falciparum in the rhoptry-neck (SURFIN4.2-RON4-GLURP) contributes to merozoite invasion. PLoS One 13, e0201669.

Rodnina, M. V, Fischer, N., Maracci, C., and Stark, H. (2017). Ribosome dynamics during decoding. Philos. Trans. R. Soc. Lond. B. Biol. Sci. 372, 20160182.

Sambrook, J. (1991). Molecular cloning: a laboratory manual (CSHL Press).

Smith, J.D., Chitnis, C.E., Craig, A.G., Roberts, D.J., Hudson-Taylor, D.E., Peterson, D.S., Pinches, R., Newbold, C.I., and Miller, L.H. (1995). Switches in expression of Plasmodium falciparum var genes correlate with changes in antigenic and cytoadherent phenotypes of infected erythrocytes. Cell 82, 101–110.

Su, X.Z., Heatwole, V.M., Wertheimer, S.P., Guinet, F., Herrfeldt, J.A., Peterson, D.S., Ravetch, J.A., and Wellems, T.E. (1995). The large diverse gene family var encodes proteins involved in cytoadherence and antigenic variation of Plasmodium falciparum-infected erythrocytes. Cell 82, 89–100.

Tonkin, C.J., van Dooren, G.G., Spurck, T.P., Struck, N.S., Good, R.T., Handman, E., Cowman, A.F., and McFadden, G.I. (2004). Localization of organellar proteins in Plasmodium falciparum using a novel set of transfection vectors and a new immunofluorescence fixation method. Mol. Biochem. Parasitol. 137, 13–21.

Trager, W., and Jensen, J.B. (1976). Human malaria parasites in continuous culture. Science (80-.). 193, 673–675.

Wahlgren, M., Goel, S., and Akhouri, R.R. (2017). Variant surface antigens of Plasmodium falciparum and their roles in severe malaria. Nat. Rev. Microbiol. 15, 479–491.

Walliker, D., Quakyi, I.A., Wellems, T.E., McCutchan, T.F., Szarfman, A., London, W.T., Corcoran, L.M., Burkot, T.R., and Carter, R. (1987). Genetic analysis of the human malaria parasite Plasmodium falciparum. Science 236, 1661–1666.

Winter, G., Kawai, S., Haeggström, M., Kaneko, O., von Euler, A., Kawazu, S., Palm, D., Fernandez, V., and Wahlgren, M. (2005). SURFIN is a polymorphic antigen expressed on Plasmodium falciparum merozoites and infected erythrocytes. J. Exp. Med. 201, 1853–1863.

World Health Organization (2018). WHO | World malaria report 2018. WHO.

Xangsayarath, P., Kaewthamasorn, M., Yahata, K., Nakazawa, S., Sattabongkot, J., Udomsangpetch, R., and Kaneko, O. (2012). Positive diversifying selection on the Plasmodium falciparum surf4.1 gene in Thailand. Trop. Med. Health 40, 79–89.

Zhang, M., Wang, C., Otto, T.D., Oberstaller, J., Liao, X., Adapa, S.R., Udenze, K., Bronner, I.F., Casandra, D., Mayho, M., et al. (2018). Uncovering the essential genes of the human malaria parasite Plasmodium falciparum by saturation mutagenesis. Science 360, eaap7847.

Zhu, X., Yahata, K., Alexandre, J.S.F., Tsuboi, T., and Kaneko, O. (2013). The N-terminal segment of Plasmodium falciparum SURFIN4.1 is required for its trafficking to the red blood cell cytosol through the endoplasmic reticulum. Parasitol. Int. 62, 215–229.

