## Supplementary material for "The pseudogene SURFIN 4.1 is vital for merozoite formation in blood stage *P. falciparum*": This file contains 4 supplementary Figures.

### Slide 1
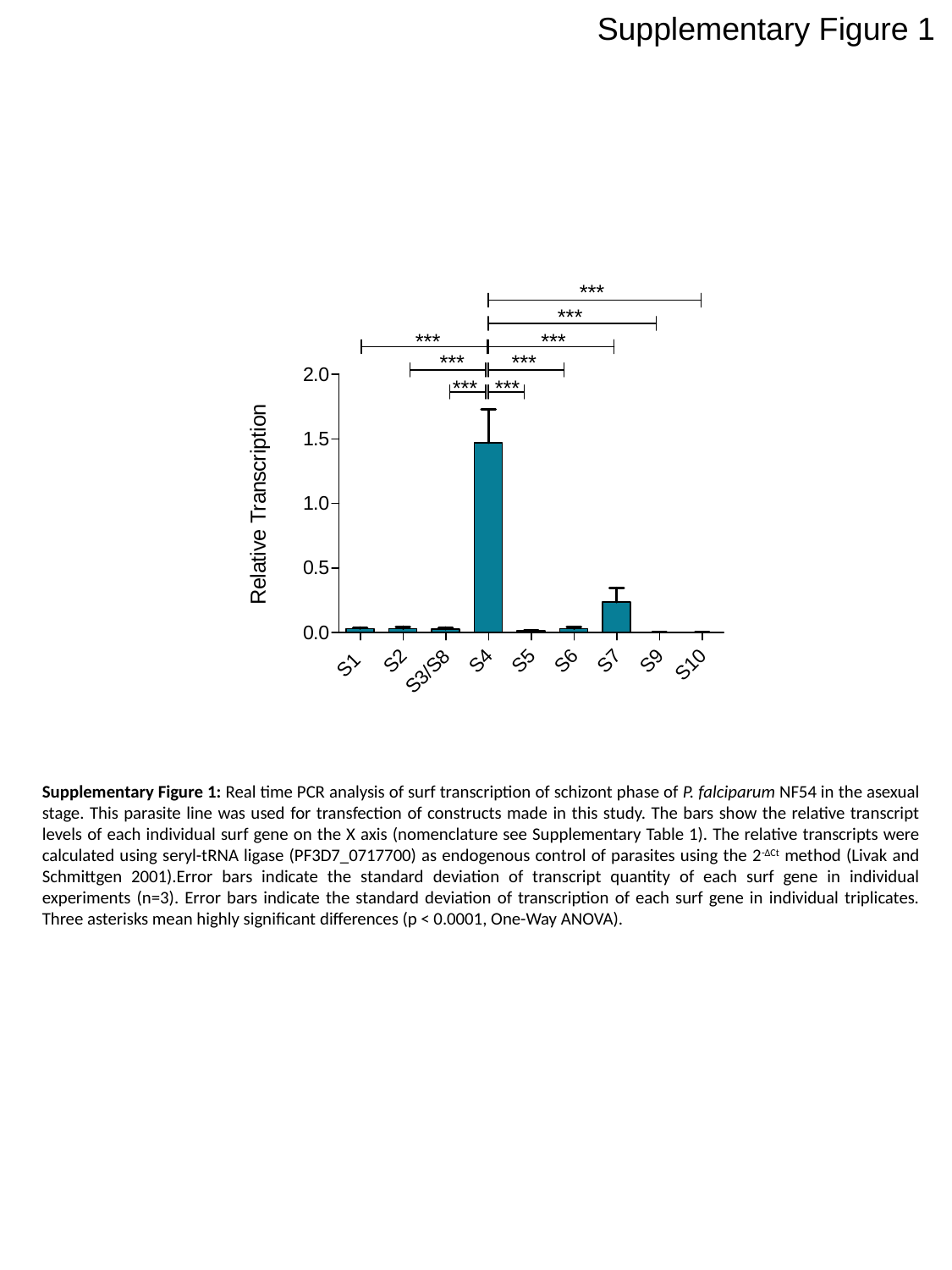

Supplementary Figure 1
Supplementary Figure 1: Real time PCR analysis of surf transcription of schizont phase of P. falciparum NF54 in the asexual stage. This parasite line was used for transfection of constructs made in this study. The bars show the relative transcript levels of each individual surf gene on the X axis (nomenclature see Supplementary Table 1). The relative transcripts were calculated using seryl-tRNA ligase (PF3D7_0717700) as endogenous control of parasites using the 2^-ΔCt^ method (Livak and Schmittgen 2001).Error bars indicate the standard deviation of transcript quantity of each surf gene in individual experiments (n=3). Error bars indicate the standard deviation of transcription of each surf gene in individual triplicates. Three asterisks mean highly significant differences (p < 0.0001, One-Way ANOVA).

### Slide 2
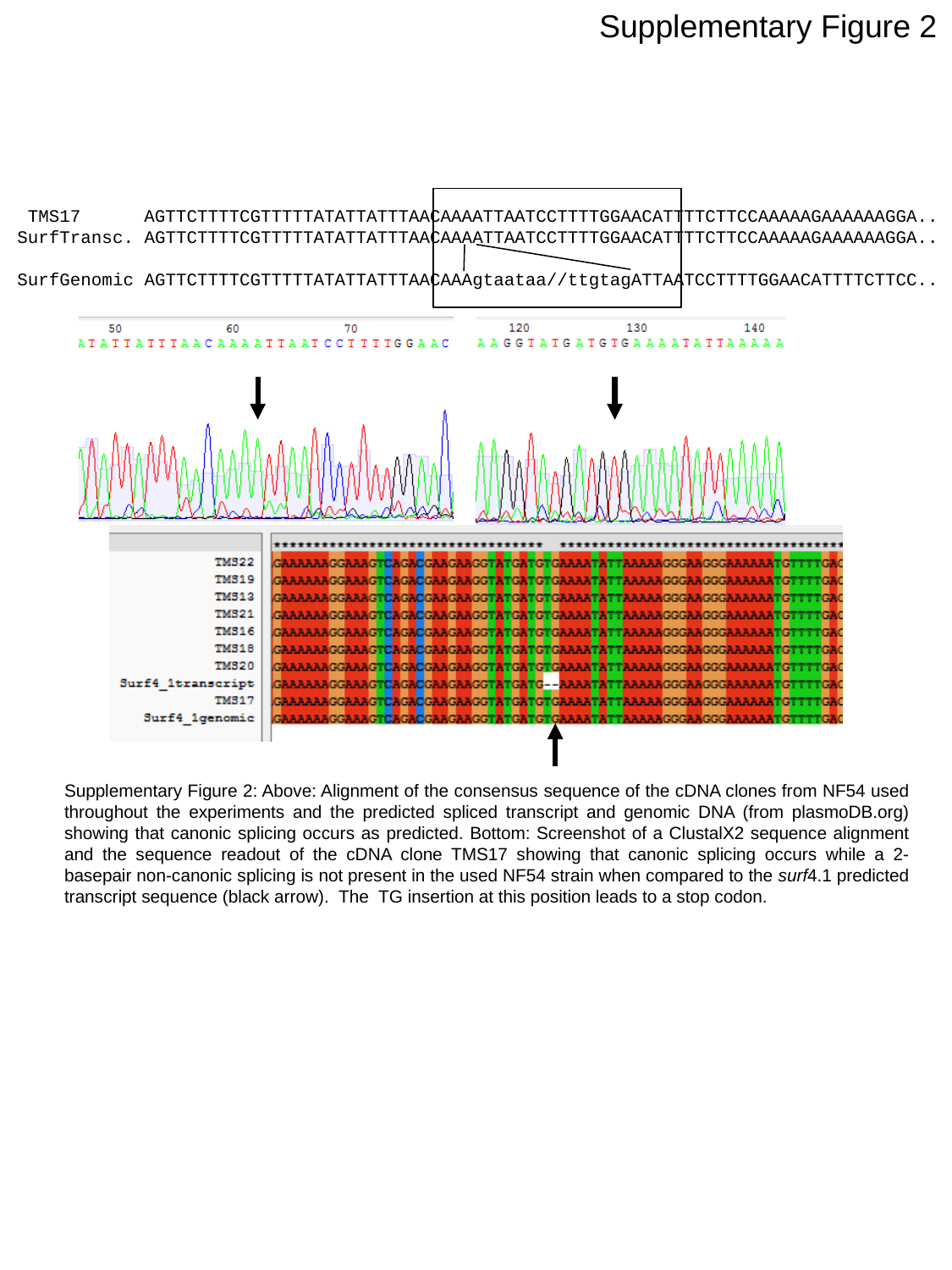

Supplementary Figure 2
 TMS17 AGTTCTTTTCGTTTTTATATTATTTAACAAAATTAATCCTTTTGGAACATTTTCTTCCAAAAAGAAAAAAGGA..
SurfTransc. AGTTCTTTTCGTTTTTATATTATTTAACAAAATTAATCCTTTTGGAACATTTTCTTCCAAAAAGAAAAAAGGA..
SurfGenomic AGTTCTTTTCGTTTTTATATTATTTAACAAAgtaataa//ttgtagATTAATCCTTTTGGAACATTTTCTTCC..
Supplementary Figure 2: Above: Alignment of the consensus sequence of the cDNA clones from NF54 used throughout the experiments and the predicted spliced transcript and genomic DNA (from plasmoDB.org) showing that canonic splicing occurs as predicted. Bottom: Screenshot of a ClustalX2 sequence alignment and the sequence readout of the cDNA clone TMS17 showing that canonic splicing occurs while a 2-basepair non-canonic splicing is not present in the used NF54 strain when compared to the surf4.1 predicted transcript sequence (black arrow). The TG insertion at this position leads to a stop codon.

### Slide 3
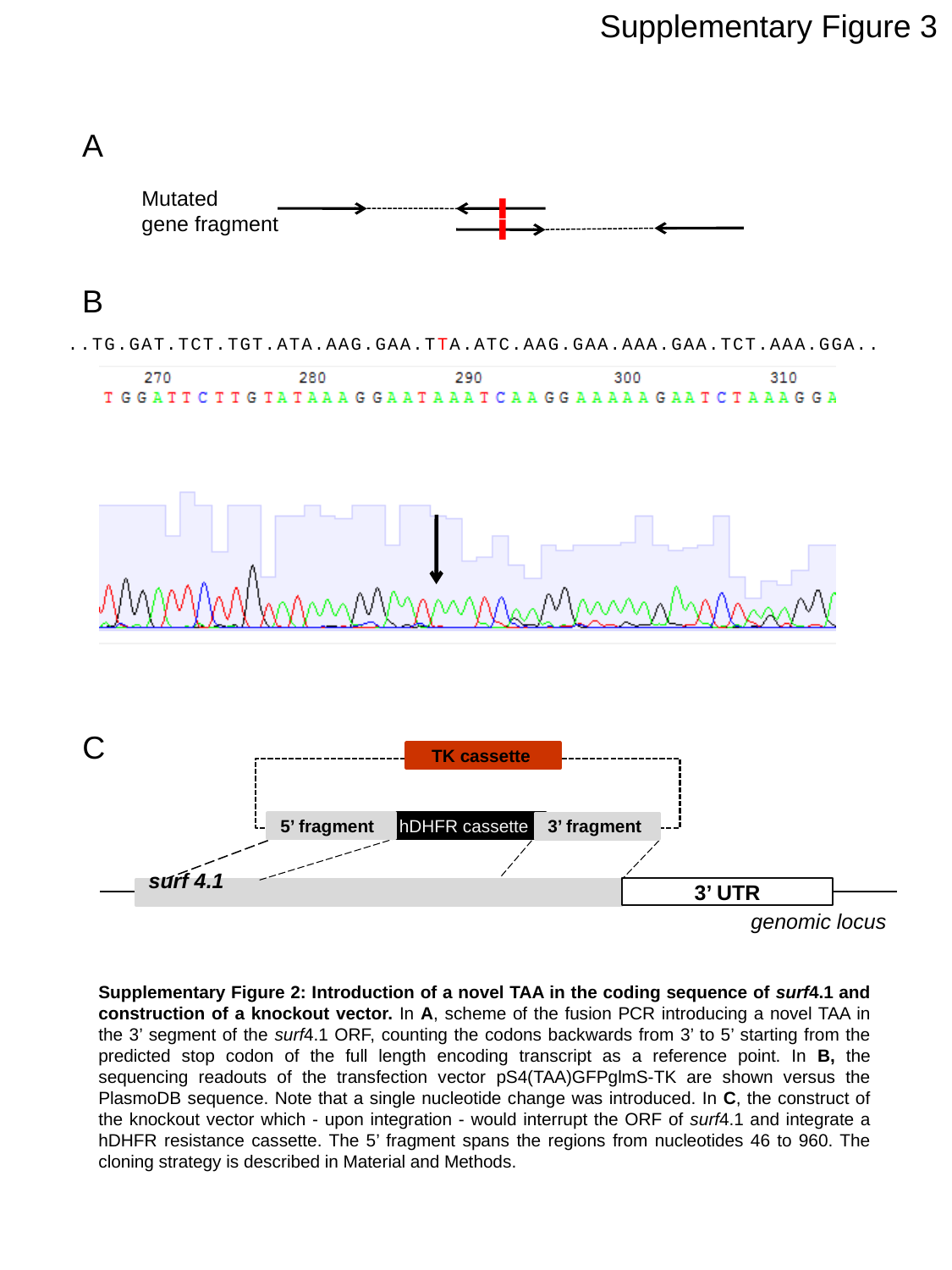

Supplementary Figure 3
A
Mutated
gene fragment
B
..TG.GAT.TCT.TGT.ATA.AAG.GAA.TTA.ATC.AAG.GAA.AAA.GAA.TCT.AAA.GGA..
C
TK cassette
5’ fragment
hDHFR cassette
3’ fragment
3’ UTR
surf 4.1
genomic locus
Supplementary Figure 2: Introduction of a novel TAA in the coding sequence of surf4.1 and construction of a knockout vector. In A, scheme of the fusion PCR introducing a novel TAA in the 3’ segment of the surf4.1 ORF, counting the codons backwards from 3’ to 5’ starting from the predicted stop codon of the full length encoding transcript as a reference point. In B, the sequencing readouts of the transfection vector pS4(TAA)GFPglmS-TK are shown versus the PlasmoDB sequence. Note that a single nucleotide change was introduced. In C, the construct of the knockout vector which - upon integration - would interrupt the ORF of surf4.1 and integrate a hDHFR resistance cassette. The 5’ fragment spans the regions from nucleotides 46 to 960. The cloning strategy is described in Material and Methods.

### Slide 4
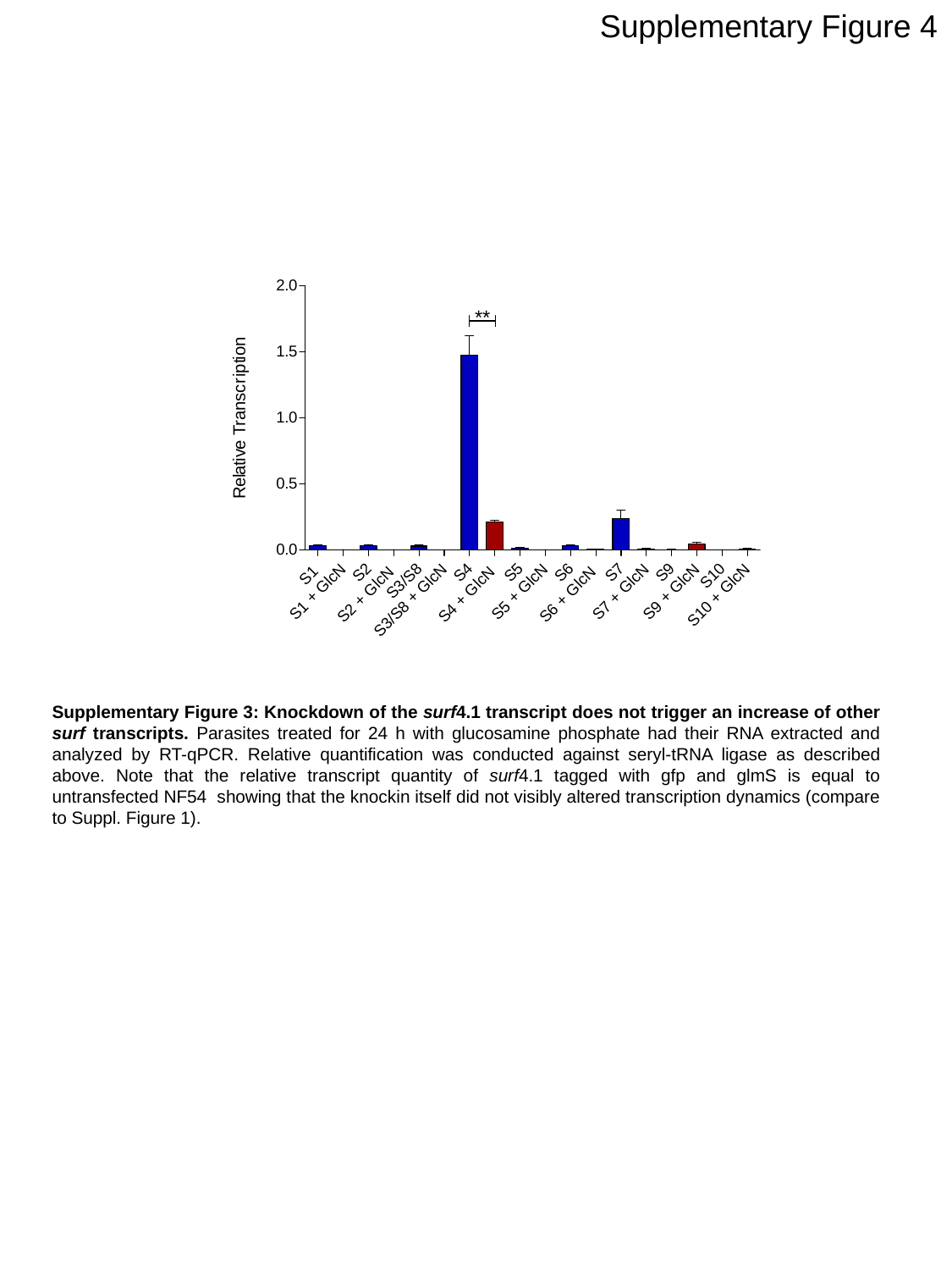

Supplementary Figure 4
Supplementary Figure 3: Knockdown of the surf4.1 transcript does not trigger an increase of other surf transcripts. Parasites treated for 24 h with glucosamine phosphate had their RNA extracted and analyzed by RT-qPCR. Relative quantification was conducted against seryl-tRNA ligase as described above. Note that the relative transcript quantity of surf4.1 tagged with gfp and glmS is equal to untransfected NF54 showing that the knockin itself did not visibly altered transcription dynamics (compare to Suppl. Figure 1).
